# PET by MRI: Glucose Imaging by ^13^C-MRS without Dynamic Nuclear Polarization by Noise Suppression through Tensor Decomposition Rank Reduction

**DOI:** 10.1101/265793

**Authors:** Jeffrey R. Brender, Shun Kishimoto, Hellmut Merkle, Galen Reed, Ralph E. Hurd, Albert P. Chen, Jan Henrik Ardenkjaer-Larsen, Jeeva Munasinghe, Keita Saito, Tomohiro Seki, Nobu Oshima, Kazu Yamamoto, Peter L. Choyke, James Mitchell, Murali C. Krishna

## Abstract

Metabolic reprogramming is one of the defining features of cancer and abnormal metabolism is associated with many other pathologies. Molecular imaging techniques capable of detecting such changes have become essential for cancer diagnosis, treatment planning, and surveillance. In particular, ^18^F-FDG (fluorodeoxyglucose) PET has emerged as an essential imaging modality for cancer because of its unique ability to detect a disturbed molecular pathway through measurements of glucose uptake. However, FDG-PET has limitations that restrict its usefulness in certain situations and the information gained is limited to glucose uptake only. ^13^C magnetic resonance spectroscopy theoretically has certain advantages over FDG-PET, but its inherent low sensitivity has restricted its use mostly to single voxel measurements. We show here a new method of imaging glucose metabolism *in vivo* that relies on a simple, but robust and efficient, post-processing procedure by the higher dimensional analog of singular value decomposition, tensor decomposition. Using this procedure, we achieve an order of magnitude increase in signal to noise in both dDNP and non-hyperpolarized non-localized experiments without sacrificing accuracy. In CSI experiments an approximately 30-fold increase was observed, enough that the glucose to lactate conversion indicative of the Warburg effect can be imaged without hyper-polarization with a time resolution of 12 s and an overall spatial resolution that compares favorably to ^18^F-FDG PET.

## Introduction

Molecular imaging seeks to characterize the fundamental molecular pathways inside organisms in a noninvasive manner. Since the rapid growth of tumors requires an abnormal metabolism to sustain it, imaging metabolism offers the possibility of detecting the transformation of tumors to a more aggressive phenotype*^1^* and of adapting treatment plans quickly in response to changes in cellular metabolic activity.*^2^* Relatively few tools exist for molecular imaging in vivo. Of these, PET using the glucose analog ^18^F-FDG is the most prominent. While ^18^F-FDG PET is an essential tool for cancer diagnosis, staging, and treatment management,*^3^* it also has its limitations. Specificity is limited in organs like the brain with a high normal glucose uptake.*^4^* Non-cancerous inflammation*^4^* and benign neoplasms*^5^* can give false positives. The background anatomical image must be taken on a different scanner, which can give rise to mis-registration errors.*^6^* Resolution in commercial PET scanners is also limited to 4-10 mm, although there are efforts to increase this limit.*^7^* The radioactivity generated by PET requires careful planning to prevent overexposure and accidental spills.*^8^* Finally, the information from ^18^F-FDG PET is limited to glucose uptake and phosphorylation, which means downstream metabolites like lactate and TCA cycle intermediates are invisible to the technique.

Glucose imaging by CEST MRI was developed to overcome some of the limitations of ^18^F-FDG PET. In glucose CEST MRI, glucose uptake is indirectly detected by saturation transfer from the exchangeable protons of glucose to water, which affords a large increase in sensitivity relative to direct detection.*^9, 10^* Due to spectral overlap, the excitation process is not entirely specific and analysis of glucose uptake is also complicated by the strong pH dependence of the exchange rate.*^11^* Also the imaging time can be quite long if adequate spatial resolution is desired.

A more direct and specific way to track *in vivo* metabolism is to follow the metabolism of exogenous tracers by magnetic resonance spectroscopy or spectroscopic imaging (MRS/MRSI). The breakdown of labeled exogenous metabolic tracers can be tracked non-invasively and precisely by ^13^C MRS. This has the advantage over static techniques, which measure the equilibrium state that may stem from many processes. However, widespread use of ^13^C MR in the clinic has been limited by poor sensitivity stemming from three fundamental reasons. First, even in the best-case scenario, the effective concentration of an exogenous contrast agent is at least 1000 times lower than that of the water signal detected in conventional MRI. Further complicating the issue, ^13^C has a low gyromagnetic ratio, which translates into a relative sensitivity only ∼2-6% that of *^1^*H for an equivalent number of nuclei. Finally, metabolic processes are inherently transient. The usual approach to overcoming low sensitivity is to simply acquire the signal for longer and average the scans. Often, however it is the metabolic flux, which is of interest rather than the steady-state concentrations, as enzymatic rates are a direct link to the activity of metabolic enzymes that are potential targets in cancer and other pathologies. Long signal averaging makes non-invasive interrogation of the rate impossible.

Dissolution dynamic nuclear polarization (dDNP) was developed to enhance signal to noise ^13^C MR and make rapid dynamic imaging of ^13^C labeled substrates and their metabolic products possible. Dynamic nuclear polarization makes use of the fact that unpaired electrons in a paramagnetic molecule can be aligned to a magnetic field to a much greater extent than the atomic nuclei with spins detected by MRI. This alignment can then be transferred to the atomic nuclei for detection. Since the signal in MRI is proportional to the degree of alignment, this polarization transfer results in a very large increase in the MRI signal, > 10,000 times in favorable circumstances.*^12^* This process occurs most efficiently at low temperatures near ∼1 K. By rapidly dissolving the frozen tracer and bringing it quickly to room temperature, a polarization sufficient to conduct metabolic MRI can be realized. After dissolution, the polarization decays by the liquid state spin lattice relaxation time of the target nuclear spin.

As a technique, dDNP is an impressive technical achievement and has demonstrated tremendous potential for metabolic imaging in vivo*^13,14^* but it has its limitations, especially in a clinical setting. Hyperpolarization is limited to a small set of molecules whose relaxation time is long enough that the enhanced polarization is not lost before the kinetics can be determined. Many key metabolites, such as glucose, have short relaxation times, and are difficult, or impossible, to image with dDNP for this reason. Since one of the fundamental limitations of ^13^C MRI is noise, it is logical that a method that reduces noise can greatly extend the utility of the technique. We show here that by considering the natural sparsity of the signal matrix, it is possible to get an order of magnitude improvement in SNR using rank reduction of the signal tensor by tensor factorization. The increase in signal to noise is large enough that it may be possible perform dynamic ^13^C tracer imaging of some ^13^C tracers without DNP.

## SVD Based Low Rank Denoising for Dynamic Single Voxel spectroscopy

We start by considering the simplest example, the dynamic non-localized pulse-acquire experiment (the same method can also be applied to dynamic localized spectroscopic imaging), as it can be described by basic concepts of linear algebra.*^15^* A key observation is that after the injection of a tracer, the intensities of peaks change as the tracer is broken down to its metabolites, but the chemical shifts are largely invariant. The time independence of the chemical shifts suggests the spectral and kinetic information are separable; an *n x m* signal matrix **M** can therefore be written as a linear combination of a small number of vectors representing spectra (**u**) multiplied by an equal number of vectors representing kinetics (**v**):*^16^*

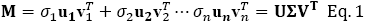

The singular value decomposition (SVD) theorem guarantees any matrix can be reconstructed fully from *n* vectors (where *n* < *m*) in this manner and the weights σ are equal to the square roots of the eigenvalues of **M^T^M**.In reality, the true signal is corrupted by noise. Noise reduction is achieved by reducing the rank of M, which is the vector space spanned by the columns (spectra) of **M**. More intuitively, the rank of **M** is the number of independent spectra *n* in Eq. 1. A rank of 1 implies the signal can be completely described as a single spectrum which decays with a single time profile. The spectra and time profile can be any arbitrary shape, but it is uniform in this case - no individual peak in the spectra decays at a different rate than any other. A signal matrix with a rank of 2 can be described as a linear combination of two spectra, with two distinct time profiles. A rank 3 matrix is a linear combination of three spectra with three distinct time profiles and so on. It is easy to see from this definition that we expect the rank (r) of the noise free matrix to be approximately equal to the number of metabolites in the sample, since noise is random and not correlated in either frequency or time. To get an approximation to **M** of rank r, we simply set the lowest *r-n* diagonal entries in **Σ** in Eq. 1 equal to zero. This result is guaranteed by the Eckart-Young-Mirsky theorem to be the best low rank approximation in the least squares sense to the original signal matrix **M**.*^17^*Since the noise-free solution is inherently low rank by the biochemistry of the problem, this solution is also likely to be an excellent approximation to the noise-free signal.

This method was first tested on data obtained from 41 mice by following the metabolism of a hyperpolarized ^13^C tracer in a single pulse (not spatially resolved) MRI experiment. A volume of 300 pL of a 98 mM solution of hyperpolarized [1-^13^C]pyruvate was injected into the tail vein of nude mice bearing tumor xenografts in the left leg. Cancer cells exhibit the Warburg effect and have higher lactate than the normal tissue. The dissolution process involved in making hyperpolarized pyruvic acid is not always perfect, resulting in spectra of varying quality. Under optimized conditions, the signal is strong enough after DNP that the main pyruvate (173 ppm) to lactate (185 ppm) conversion can be easily quantified without additional signal processing. In others, the signal is barely detectable. Since the biochemistry is the same in each of these cases, we know *a priori* the peak positions and approximate kinetics in the noisy data. This property makes the pyruvate dDNP experiment an excellent test platform for the accuracy of signal reconstruction.

## SVD Based Low Rank Denoising Gives an Order of Magnitude Improvement in SNR in DNP ^13^C Tracer Experiments

**Figure 1A** shows an example of noisy pyruvate and lactate peaks in a dDNP experiment. The smaller peaks corresponding to alanine and pyruvate hydrate are completely buried within the noise. Even averaging 5 scans together is insufficient to accurately quantify the lactate or to detect the minor peaks. Averaging also results in a significant loss of time resolution **(Figure 1A)**. Rank reduction by SVD gives a substantial (9.3 fold) improvement in signal to noise **(Figure 1C)**. If we define signal to noise as the maximum value of the most intense peak divided by the standard deviation of the signal in a region of the where no peak is expected, signal to noise increased by nearly an order of magnitude (mean=9.4, median=7.3) after rank reduction to a rank of 5 **(Figure 1D).**

**Figure 1.**
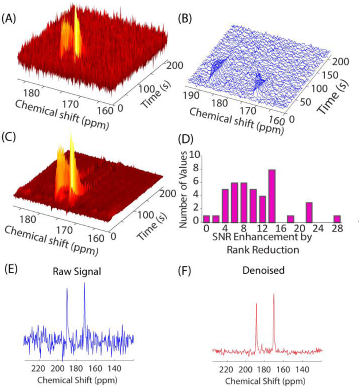
SNR improvement in noisy data using rank reduction. **(A)** Dynamic data from a 13C pyruvate tracer single pulse dynamic nuclear polarization experiment with low signal to noise. **(B)** Signal from A averaged over 5 scans. Even with averaging, the signal is too weak to be quantified accurately. **(C)** Signal reconstruction using rank reduction to a rank of 5. The peaks corresponding to the two main metabolic products pyruvate and lactate are now clearly visible in the reconstructed spectra, along with two minor peaks corresponding to alanine and pyruvate hydrate side products. **(D)** Histogram of the SNR improvement using rank reduction (r=3) over 41 mice. SNR is defined here as the intensity of the maximum signal divided by the standard deviation in a 40_point region of the spectrum where signal is known not to be present. **(E and F)** Slice from **A** and **C** showing the 9 fold SNR enhancement after reconstruction.

## Previously Undetectable Peaks Can be Quantified by Low Rank Denoising

This level of noise reduction enables some measurements that were previously considered difficult. The bicarbonate provides information on the balance between the glycolytic and oxidative metabolic pathways*^18,19^* and in combination with the CO2 signal can serve as a measure of intracellular pH.*^20^* The bicarbonate signal is difficult to detect in the raw spectra even in hyperpolarized experiments **(Figure 2A and C**)*^18^*. After rank reduction by SVD, the bicarbonate signal becomes evident **(Figure 2B and D)** and breakdown of pyruvate through the citric acid cycle in the oxidative phosphorylation pathway can be followed (**Figure 2F)**. The detection of the bicarbonate peak at the expected position shows that rank reduction in the kinetic domain gives an actual increase in sensitivity, as opposed to the cosmetic improvement that eliminates both noise and true weak signals from some other noise suppression techniques.*^21^*

**Figure 2.**
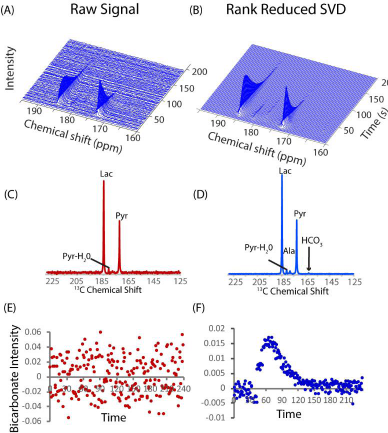
Detection and quantification of weak peaks using rank reduction Left. Raw data from a pyruvic acid DNP experiment from Figure 1 with high signal to noise. Pyruvate (173 ppm) is converted to lactate (185 ppm). Only the pyruvate, lactate, and pyruvate hydrate signals are quantifiable. Right Rank 5 reconstruction of the raw data. In the raw data (**A** and **C**), the bicarbonate peak at 162.5 ppm is not detectable even though the overall signal to noise is high. Using the low rank approximation **(B and D)** a kinetic profile of bicarbonate metabolism can be obtained **(F)**. No prior information about the bicarbonate signal or kinetics was used.

## Higher SNR Translates to More Precise Kinetic Fitting

The performance seen in **Figures 1** and **2** may not be universal as noise reduction algorithms are often sensitive to the characteristics of the signal.In particular, large dynamic ranges are often problematic for many noise reduction algorithms. **^21^**To confirm the generality of the method, we also tested the method using hyperpolarized [2- ^13^C]pyruvate. C-l labeled pyruvate restricts analysis to the first steps of the TCA cycle as the ^13^C label is released as ^13^C-bicarbonate. Hyperpolarized [2- ^13^C]pyruvate allows tracking further downstream in the TCA cycle and into other metabolic pathways.*^22,23^* The data using hyperpolarized [2-^13^C]pyruvate shows a dominant C-2 pyruvate peak at 208 ppm and weak peaks near the noise level corresponding to the downstream metabolites 5-glutamate and 1-acetyl-carnitine at 184 ppm and 175 ppm, among other peaks (**Figure S1B**). The downstream metabolite peaks are less than 1% of the substrate signal. Rank reduction to a rank of 5 independent species substantially improves the signal to noise of later time points where the weaker downstream metabolite peaks are no longer visible in the raw signal (compare **Figure S1C** to **S1E**). The signal to noise increase translates to a corresponding increase in the accuracy of the kinetic reconstruction. The 8.3-fold increase in SNR translates to a 50 fold increase in the precision of the influx rate *ki* corresponding to the conversion of pyruvate to glutamate through the TCA cycle and a 200 fold increase in the precision of the loss rate *k2* corresponding to signal loss through relaxation and transformation to other metabolites. (compare **Figure S1H** to **S1G**). The same data was also tested with another state of the art denoising technique, wavelet shrinkage. While wavelet shrinkage using soft thresholding on individual spectra was unsuccessful in significantly reducing the noise (**Figure S1D**), SVD rank reduction (**Figure S1E**) was able to detect and quantify the weak peaks by using the additional information present in the entire time course.

## Denoising Does Not Bias Relative Intensities or Kinetics

Rank reduction could introduce bias into curve-fitting of the kinetics, limiting its usefulness as a noise reduction technique. One method to test bias is to simulate realistic spectra and time courses and test the accuracy of the method by kinetic modeling in the presence of increasing amounts of noise. The pyruvate DNP experiment of Figure 1 is used as a test case. Nonlinear regression is sensitive to small levels of noise and curve fitting was imprecise with even modest noise levels (**Figure S2**). Since potential bias was difficult to measure with curvefitting when the measurements are so imprecise, we used an alternate technique, the Area Under the Curve (AUC) approach, which is more robust against noise as the kinetic constants are derived from a ratiometric sum over all time points. Any potential bias introduced by SVD can therefore be measured at high levels of noise by this method even when traditional curve-fitting fails. The drawback is that transporter uptake cannot be quantified and the method is difficult to apply to models more complicated than three site exchange.

The accuracy of SVD reconstruction depends on the SNR of the metabolite peak. When the noise level is significantly less than the metabolite peak (SNR of the metabolite peak >10), no bias is introduced by SVD; the kinetic constants of all metabolites can be recovered accurately by the AUC method when SVD is used with a rank of 5 or higher (**Figure S2**). Since the SNR of the pyruvate and lactate peaks is almost always above 10 in DNP experiments, any effect of SVD on these metabolites will be minimal in most cases. The kinetics of bicarbonate can also be recovered without bias at all levels of noise **(Figure S3A**). At higher levels of noise; however, there is a slight tendency for the alanine signal to drift towards the kinetics of the stronger pyruvate signal (**Figure S3B**). The pyruvate to alanine conversion rate is overestimated by ∼ 10% when SVD reconstruction. The origin of this error is that at low SNR the 4^th^ and 5^th^ kinetic eigenvectors that differentiate the kinetics of alanine from the other signals become indistinguishable from noise. While this suggests the error could be corrected by using a larger rank in the reconstruction, adding more eigenvectors decreases the precision to the point that the method becomes ineffective.However, the effect is modest and confined to the alanine signal under ordinary conditions. Overall, the simulations suggest rank reduction by SVD does not introduce a significant bias into measurements of either kinetics or intensities in DNP experiments under normal conditions.

Synthetic data is less reliable for testing denoising algorithms than actual data due to the assumed idealities in the simulation. Noise in real data may show a frequency or time dependence that may differ from an ideal Gaussian white noise distribution,*^24^* and kinetics often do not exactly follow simple models. To test the accuracy of rank reduction on a more realistic sample, we performed a ^13^C MRI DNP experiment using hyperpolarized [1- ^13^C]pyruvate, with excitation pulses alternating between a 10^°^ pulse on odd scans and 2^°^ pulses on even scans to generate high and low noise datasets from the same sample (**Figure 3**). SVD reduction of the low flip angle signal to a rank of five gave a 3-fold increase in signal to noise. The kinetic profiles of the reconstructed pyruvate, pyruvate hydrate, and lactate signals, which have relatively strong signals that could be accurately measured even at low flip angles, closely match the high flip angle signal when normalized for intensity (**Figure 3B-D**). The reconstructed signal of alanine also nearly exactly matches the high-power signal despite the significant noise in the low power signal (**Figure 3E**). Only the very weak bicarbonate signal, which is completely unrecognizable in the raw signal (**Figure 3F**), shows a slight error in reconstruction.

**Figure 3.**
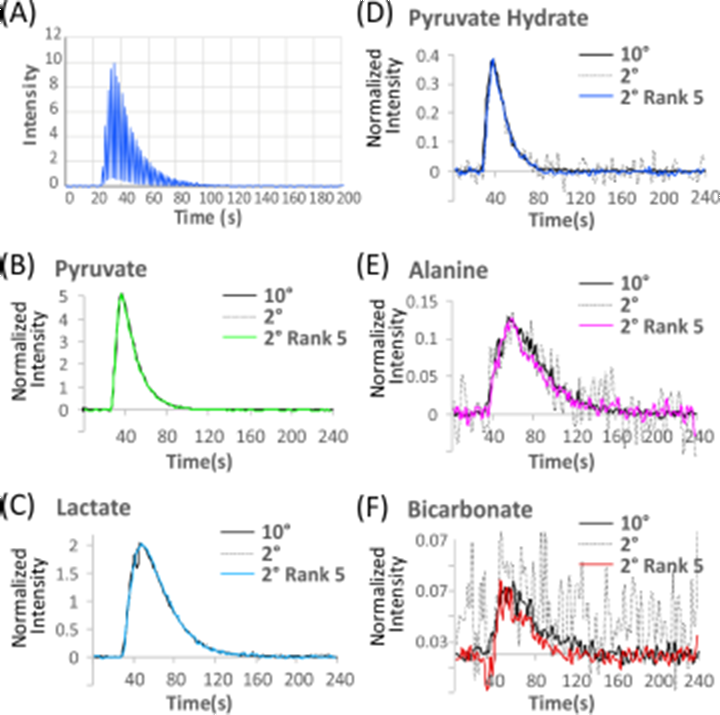
Accuracy of low rank SVD reconstruction. (A) Pyruvate signal after the injection of hyperpolarized 1- C pyruvic acid using 2° or 10° flip angles on the odd and even pulses respectively. Even with a reduction to one fifth the original signal the kinetic profile of pyruvate (B), lactate (C), pyruvate hydrate (D), and alanine (E) are reconstructed almost exactly by SVD (colored lines). The bicarbonate signal (F) contains only a minor error in the breakdown constant, despite the pyruvate hydrate signal being almost undetectable in the raw data (grey dots).

## Dynamic Single Voxel Spectroscopy of Glucose Metabolism without DNP

The results from DNP experiments encouraged us to try the SVD processing on molecules less amenable to hyperpolarization than pyruvate. Increased glucose uptake to meet the increased energetic and synthetic demands of increased cell growth is common feature of many cancers*^25,26^*, which makes glucose a primary target for metabolic imaging through both PET*^27^*and MRI techniques like CEST.*^25,29^* Unfortunately, the short T1 of glucose causes rapid relaxation of the hyperpolarized signal, meaning most of the signal is lost before ^13^C glucose enters the glycolytic pathway.*^30^* **Figure S4** shows the results from a tail vein injection of 50 pL of 100 mM uniformly ^13^C labeled glucose, deuterated at all non-exchangeable protons to increase T1 (see **Table S1**), into a mouse with a leg xenograft from the Hs766t metastatic pancreatic carcinoma cell line. Even with deuteration, most of the hyperpolarized signal is lost within the first ten seconds and unavailable for detection of downstream metabolic products (**Figure S4**). Nothing besides the glucose peak can be detected in the raw signal (**Figure S4**). In the signal rank reduced by SVD; however, a very faint signal of approximately 0.5% the intensity of the glucose peak can be seen at 184 ppm corresponding to the 1 - position of lactate. Like the glutamate and bicarbonate signals before, SVD brings the weak lactate signal up to detectability.

With hyperpolarized glucose, the DNP scans were of limited value due to the rapid signal decay. The success of SVD rank reduction in suppressing noise encouraged us to image uniformly labeled glucose solutions without hyperpolarization. The first studies were non-localized spectroscopy experiments similar to the ones described above. **Figure 4** shows the results before and after SVD rank reduction from a tail vein injection of 350 pL of 555 mM uniformly ^13^C labeled glucose into a mouse with a leg xenograft from the Hs766t cell line taken on a 9.4 T scanner (see Materials and Methods for details). The scans show a gradual uptake of glucose and a subsequent breakdown to lactate and alanine, reflecting transport limited uptake and the primarily aerobic glycolysis in this particular cell line*.*^31^** Every peak of both the α- and β anomers of glucose were identified and quantified (**Figure 4D-F)**. Glucose-6-phosphate was identified as a shoulder of the 6’ carbon of glucose with distinctly different kinetics*^32^* (**Figure 4E-F**). In addition to alanine, lactate, glucose, and glucose, glucose-6-phosphate, a weak peak belonging to glycogen at 99 ppm**^32^** was identified in some of the samples with high signal to noise. As a control, the experiment was repeated in a mouse without a tumor xenograft. No glucose peaks could be seen in the absence of a tumor (**Figure S5**), confirming the signal originates from the enhanced uptake and retention of glucose in tumors. The overall time-scale of glucose metabolism approximately matched previous ^13^C measurements but with greatly increased time resolution.*^33^* Previous measurements were limited by signal-to-noise to taking one spectra every 5 minutes, while a quantifiable signal can be acquired using rank reduction every 3.2 s. The increased temporal resolution allowed measurement of the rate of uptake of glucose, which was challenging with the previous ^13^C measurements, and difficult even with PET imaging.*^34 35^* The experiments in **Figure 4** were made on a 9.4 T scanner, which is not clinically widely available. With an eye towards eventual clinical translation, we made corresponding measurements at 3T (**Figure S6**). Although the lactate signal was not resolved, possibly because decoupling was not implemented, both glucose uptake and metabolism could be clearly quantified.

**Figure 4.**
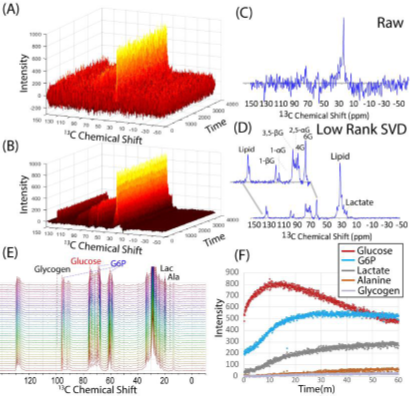
Dynamic Single Voxel Spectroscopy of Glucose Metabolism without DNP. (A) Raw 13C signal after injecting 50 mg of uniformly 13C labeled glucose into the tail vein of a mouse with a MiaPaca xenograft without hyperpolarization. (B) The same signal after rank reduction by SVD to a rank 5. (C) Raw 13C signal after 1000 seconds. (D) The same signal as (C) after rank reduction by SVD. While only the lipid signal could be detected clearly in the raw signal, every peak of both the α- and β anomers of glucose can be identified and quantified in rank reduced signal. Peaks belonging to lactate, alanine, and glucose 6-phosphate can also be identified in the rank reduced signal. (E) Stacked plot showing the evolution of the glucose signal in an experiment with high SNR. Glucose-6- Phosphate (G6P) can be identified as a growing shoulder along some of the glucose peaks. (G) Kinetics of the metabolite peaks identified from (D)

## Extending SVD into Higher Dimensions through Tensor Decomposition

SVD is a strictly two-dimensional matrix method and cannot be used directly on images of this type. In order to adapt this method to higher dimensional data typical of dynamic, volumetric, and spectroscopic medical imaging experiments, a different method is proposed. The individual voxels can be treated independently and denoised by SVD but the fewer time points are acquired for each voxel in an imaging experiment. Because noise reduction power of SVD varies with the matrix size by Q/r, where Q is the smallest matrix dimension and r is the predicted rank,36 using SVD to denoise individual voxels is not powerful enough for imaging experiments (see below).

Treating voxels independently ignores the correlation between voxels and the low rank structure of the overall image37. A multidimensional analog of the SVD, the Tucker Decomposition, can be used to find a low rank reconstruction of the entire image. Similar to the SVD, the Tucker Decomposition factorizes an n-dimensional tensor X into a core tensor G of the same dimensions as the original data array and a set of n factor matrices {A,B,C,…}:*^38^*

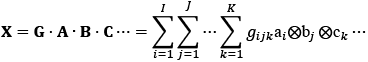

Where ⊗ refers to the tensor outer product. While SVD decomposes a data matrix into a linear combination of *vectors,* the Tucker Decomposition decomposes a higher dimensional data tensor into a multilinear combination of *matrices* corresponding to images and time-dependent spectra (see **Figure 5**). Like the SVD, the core tensor can be truncated to suppress noise while retaining as much of the signal as possible. The use of a more complex basis allows the underlying structure of the data to be represented in a more compact and natural way, which translates to more effective denoising when the distribution of signals is not random.*^39^*

**Figure 5:**
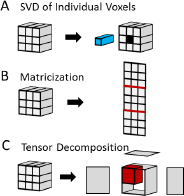
Methods of data reduction for a data tensor. For simplicity, the data tensor is shown here as a three dimensional object. The actual data tensor in the dynamic CSI experiment is four dimensional. (A) SVD of individual voxels. Each voxel is treated independently and rank reduced by SVD. (B) Matricization. The *I Χ J Χ K Χ L* four dimensional data tensor is unfolded into a two dimensional (I+J+K) Χ L matrix. The resulting matrix is then rank reduced by SVD and then refolded to the original dimensions of the data tensor (C) Tensor Decomposition. The *I ΧJ Χ K Χ* L data tensor is factorized into a set of factor matrices multiplied by a sparse tensor with the same dimensions as the original data set. Rank reduction is obtained by truncating the values along each dimension.

## Dynamic CSI Imaging of Glucose Metabolism without DNP by Tensor Decomposition Rank Reduction

**Figure 6** shows the results from dynamic chemical shift imaging of a 50 mg bolus of [U-^13^C]glucose injected into the tail vein of a mouse with a leg xenograft from the MiaPaca cell taken at 9.4 T before and after tensor decomposition. One image was taken every 48 s. The raw image is predominantly noise **(Figure 6C**) with no discernable signal detectable in the voxel (**Figure 6E** and **F**). A very weak glucose signal can be detected in some voxels by using voxel by voxel SVD with a high threshold for rank reduction (**Figure 6G** and **H**), a roughly 3-fold improvement in SNR compared to the raw signal (see **Figure 6M)**. More aggressive rank reduction distorts the image by suppressing the signal in some of the voxels. The multi-dimensional nature of the tensor decomposition is essential to its success, unfolding the data into a 2D matrix and then using SVD low rank reconstruction (similar to the initial step of the LORA technique (see methods)*^40^*, **Figure 5B**) resulted in a 3-fold less improvement in SNR on this data set (**Figure 6I and J**). Only by tensor factorization is a clear glucose signal detectable in each of the time traces, giving on average a ∼31 fold improvement in SNR (**Figure 6M)**. The improvement in SNR is large enough that the SNR for each time point is actually higher than the SNR obtained by averaging over the entire time course of 90 min (450 scans) and sufficient to image the lactate signal in addition to the lipid and glucose signals. The time traces from the CSI images show relatively rapid uptake of glucose within the tumor that reaches a plateau within approximately 10 min, a similar time scale as the non-localized experiment. Glucose uptake is almost entirely localized within the tumor with a clear separation from lipid signal originating from the gluteal fat pad in the upper leg **(Figure 6A)**. ^13^C lactate builds up slowly as glucose is broken down **(Figure 6C and D)**. To our knowledge, this represents the first successful dynamic imaging of tracer metabolism *in vivo* through ^13^C MRSI without dynamic nuclear polarization.*^30^*

**Figure 6:**
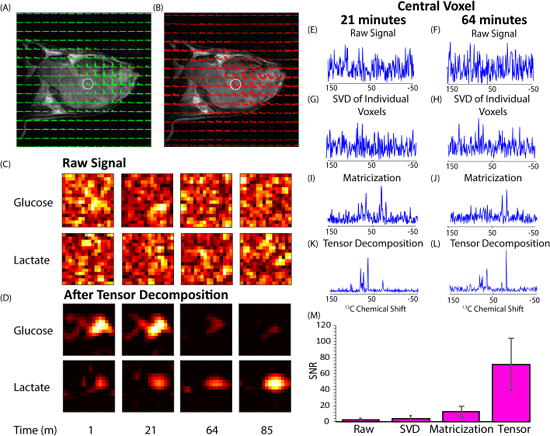
Dynamic C CSI Imaging of Glucose Metabolism without DNP. CSI imaging of a mouse leg Hs766t xenograft after a 50 mg glucose injection. An 8×8 image of the tumor bearing mouse leg was acquired by chemical shift imaging every 48 seconds for 90 minutes. The final image was zero-filled to 16×16. Each is voxel 0.15 cm x 0.15 cm x 1.6 cm in size. **(A)** The glucose region of the spectra for scan 44 overlaid on the anatomical image after tensor_3_ decomposition. **(B)** The kinetics. **(C and D**) Contour maps created from the peak maximums of the glucose and lactate C signals at the time points indicated. While the raw images are uninterpretable, the images after tensor decomposition closely conforms to the boundaries of the tumor and a clear difference in the between the kinetics of glucose and lactate can be detected. **Effect of signal processing on SNR (E to L)** The spectrum of central voxel at the indicated time points using different processing techniques. **(E and F)** No peaks are evident in the raw signal. **(G and H)** Using SVD on each voxel independently slightly improves SNR, enough to detect a very weak glucose signal **(I and J)** Introducing global correlations by matricizing the data tensor and using a single SVD simultaneously on all the data yields a detectable but still noisy glucose signal. **(K and L)** Tensor decomposition allows a more natural representation of the dour dimensional signal, which translates into greatly improved signal to noise (SNR=27). **(M)** Histogram of the SNR with each technique over 14 mice. Tensor factorization yields a 31 fold improvement in SNR, approximately 3 times better than matricization.

## Noise reduction without a spectral dimension

Spectroscopic imaging is difficult in hyperpolarization experiments due to the irreversible loss of hyperpolarization, which sets a limit on how many scans can be acquired before the signal becomes undetectable. One method of rapidly acquiring a metabolite specific image is to use a fast imaging sequence like EPI in combination with spectrally selective pulses switching the excitation profile on alternate scans to center on different metabolites. The result is a series of images that correspond to the distribution of the metabolite in time.

To test the usefulness of the low rank approximation on these types of experiments, as well as other dynamic MRI experiments where spectral information is not available, we tested low rank reconstruction on spectrally selective EPI images of lactate metabolism after the delivery of a bolus of hyperpolarized ^13^C-1-pyruvate. In the raw data, only the kidneys and part of the liver can be seen in the individual time points making the goal of following pyruvate metabolism at the organism level difficult. Using tensor reconstruction, the signal to noise is improved by a factor of 3 (**Figure S6**). With the increase in signal to noise, the heart became visible and the liver and kidneys were more clearly defined. With the reconstructed model, it is possible to trace the production of lactate through the body, detected in the heart by frame 3 and being retained in the liver and kidneys until frame 12. While the increase in signal to noise is less impressive than in spectral images, it does show that tensor factorization may be a viable technique for increasing the SNR in low signal, low complexity images.

## Discussion

The possibility of using ^13^C MRI for molecular imaging was investigated early on in the history of MRI but is not used routinely once it became apparent that the SNR was insufficient for imaging. If this limitation can be overcome, ^13^C MRI with uniformly labeled ^13^C glucose may become an alternative/complementary technique to ^18^F-FDG PET. We show here that this limitation is not as formidable as may have once seemed. The key to this development is a relatively simple but efficient and robust post-processing noise suppression technique that takes advantage of the common underlying structure in metabolic imaging experiments, particularly those with a kinetic component. There have been a number of attempts at denoising individual magnetic resonance spectra, either specifically in the context of MRSI or more generally throughout the NMR field. The methods can be divided into those that attempt to denoise the free induction decay (FID) signal in the time domain (Cadzow reduction*^41,42^* aka HLSVD*^42,43^*, HTLS*^44,46^* and Pade transform methods) and those that attempt to directly denoise the MR spectrum in the frequency domain. Time domain methods make the assumption that the signal originates from a small set of ideal Lorentzian peaks with well-defined frequencies, intensities, and line widths.*^47^* From this assumption, a variant of the rank reduction through SVD procedure is employed to distinguish true peaks from false noise peaks, using a sliding window to convert the FID signal into a matrix. In ideal conditions, this results in a perfect, noise-free spectrum. However, the true rank of the noise-free signal is difficult to estimate *a priori* from the FID. Imperfect shimming, magnetic susceptibility artifacts, or chemical exchange processes can cause a deviation from ideal line-shapes. Any peak with a non-ideal line-shape manifests as multiple Lorentzian peaks in the fitting procedure. For example, the residual water peak in *^1^*H MR can only be modeled by at least 4 Lorentzian peaks.*^45^* In many cases, weak peaks like the bicarbonate peak in **Figure 2** are suppressed by this procedure in the presence of strong peaks since most of the degrees of freedom are used in modeling more accurately the line-shape unless the rank is set to a fairly large number, which has the undesirable consequence of creating spurious peaks. Quantitation may also be difficult in many cases unless the spectrum is largely noise free.*^49^* To surmount these problems, one variant of the LORA technique *^49-55^* uses a low rank approximation in the spatial domain before attempting to denoise the FID of individual spectra. The method is not perfect in suppressing artifact peaks and necessarily results in a loss of spatial resolution.

Other methods denoise the MR spectrum directly. The total variation approach uses a smoothing operator to dampen sharp discontinuities that likely reflect noise.*^56^* Wavelet methods seek to decompose the signal into subsignals of increasing complexity, with the highest complexity signals, which likely represent noise, thrown out.*^57,58^* Maximum entropy methods take a similar approach form the perspective of information theory. These methods have the disadvantage of low sensitivity in the sense that they tend to flatten weak peaks*^21^* and may eliminate other fine details of the spectra.

Despite their differences, all of these methods operate in the level of individual spectra. By considering rank reduction in the kinetic domain specifically, where the signal matrix is particularly sparse and possesses a natural connection to the underlying chemistry, it is possible to achieve an order of magnitude or more improvements in SNR without sacrificing resolution or accuracy. This level of enhancement can detect dynamic signals in the presence of large amounts of noise (**Figure 1**) and quantify weak signals even in the presence of much stronger signals **(Figure 2**). For dDNP experiments, this opens up the possibility of using lower flip angles and doses than are currently being used. Since the transverse magnetization after the i^th^ pulse in dDNP depends on the flip angle α according to sin(a)cos^1-1^(a), improvements in SNR coupled with low flip angles may enable the detection of metabolites further downstream than is currently possible. Higher signal-to-noise is also useful in expanding the range of chemical probes amenable to DNP. Substrates with low polarization efficiencies or short T1 relaxation times like glucose (**Figures 4 and 6**) can be quantified more easily using SVD or tensor factorization. Finally, higher SNR may open up the possibility of off-site hyperpolarization through the brute force approach,*^59^*’ **^60^** which would eliminate the largest barrier for clinical use.

Some challenges associated with the translation of ^13^C glucose imaging into a clinical technique remain to be overcome. The 2g/kg dose used in this study is about three times larger than the maximum tolerated IV dose. **^61^**Whole body imaging as done in ^18^F-FDG PET/CT will be difficult with the technique. To get adequate resolution in humans, the number of k-space points will need to be larger, around 64×64 for a brain scan with a FOV of approximately 190 mm and 3 mm voxels, for example. This will result in unacceptably long acquisition time for each scan if conventional CSI imaging is used since each k-space point is phase encoded and requires a separate RF excitation and data acquisition. These are not insurmountable difficulties and could possibly be addressed with a faster imaging sequence, improved *^1^*H decoupling, and parallel imaging.

## Conclusions

### Materials and Methods

#### Mouse Models

The animal experiments were conducted according to a protocol approved by the Animal Research Advisory Committee of the NIH (RBB-159-2SA) in accordance with the National Institutes of Health Guidelines for Animal Research. Female athymic nude mice weighing approximately 26 g were supplied by the Frederick Cancer Research Center, Animal Production (Frederick, MD) and housed with *ad libitum* access to NIH Rodent Diet #31 Open Formula (Envigo) and water on a 12-hour light/dark cycle. Xenografts were generated by the subcutaneous injection of 3 ×10*^6^* MiaPaCa-2 (America Type Cell Collection (ATCC), Manassas, VA, USA) or Hs766t (Threshold Pharmaceuticals, Redwood City, CA, USA) pancreatic ductal adenocarcinoma cells. Both cell lines were tested in May 2013 and authenticated by IDEXX RADIL (Columbia, MO) using a panel of microsatellite markers

#### 13C MRS with Dynamic Nuclear Polarization

[1-^13^C]pyruvic acid (30 μL), containing 15 mM TAM and 2.5 mM gadolinium chelate ProHance (Bracco Diagnostics, Milano, Italy), was hyperpolarized at 3.35 T and 1.4 K using the Hypersense DNP polarizer (Oxford Instruments, Abingdon, UK) according to the manufacturer’s instructions. Typical polarization efficiencies were around 20 %. After 40-60 min, the hyperpolarized sample was rapidly dissolved in 4.5 mL of a superheated HEPES based alkaline buffer. The dissolution buffer was neutralized with NaOH to pH 7.4. The hyperpolarized [1- ^13^C]pyruvate solution (96 mM) was intravenously injected through a catheter placed in the tail vein of the mouse (12 μL/g body weight). Hyperpolarized ^13^C MRI studies were performed on a 3 T scanner (MR Solutions, Guildford, UK) using a home-built ^13^C solenoid leg coil. After the rapid injection of hyperpolarized [1-^13^C]pyruvate,spectra were acquired every second for 240 s using a single pulse acquire sequence with a sweep width of 3.3 kHz and 256 FID points.

#### Dynamic ^13^C Glucose MRS without DNP

Magnetic resonance spectroscopy was performed on either a 9.4 T Biospec 94/30 horizontal scanner or a MR Solutions 3 T horizontal scanner. The coil assembly for the mouse leg consists of 3 16 mm independent wire loops that are each terminated with a double balanced tune/match network to the 50 Ohm characteristic impedance of the coaxial cable. The two ^13^C coils are geometrically decoupled because they have their radiofrequency field orthogonally oriented. Those coils are fed with radiofrequency currents with a 90° phase difference. One of the coils has a solenoidal shape and the other is constructed by two saddle loops with 120° arcs (quasi-Helmholtz pair), arranged coaxially to the solenoidal coil. The ^1^H coil is a double sized surface coil coaxially arranged to the solenoidal coil. The proton coil has integrated ^13^C frequency traps and the ^13^C coils have integrated *^1^*H frequency traps to minimize coupling between them. A bandpass filter was used to minimize contamination of the ^13^C signal by the *^1^*H decoupling pulse.

Each mouse was anesthetized during imaging with isoflurane 1.5-2.0% administered as a gaseous mixture of 70% N2 and 30% and kept warm using a circulating hot water bath. Both respiration and temperature were monitored continuously through the experiment and the degree of anesthesia adjusted to keep respiration and body temperature within a normal physiological range of 35-37° C and 60-90 breaths per min. Anatomical images were acquired with a RARE fast spin echo sequence*^62^* with 15 256×256 slices of 24 mm × 24 mmx 1 mm size with 8 echoes per acquisition, a 3 s repetition time, and an effective sweep width of 50,000 Hz. Samples were shimmed to 20 Hz on the 9.4 T with first and second order shims using the FASTMAP procedure.*^63^* Non-localized spectra of glucose without DNP at 9.4 T were acquired with the NSPECT pulse-acquire sequence using maximum receiver gain, a repetition time of 50 ms, Ernst Angle excitation of 12°, 256 FID points, a sweep width of 198.6 ppm, 16 averages per scan, and 4500 scans for a total acquisition time of 1 hour. MLEV16 decoupling*^64,65^* was applied during acquisition using −20 dB of decoupling power and a 0.2 ms decoupling element. The decoupling pulse was centered on the main proton lipid resonance at 1.3 ppm. Data at 3 T was acquired similarly except decoupling could not be applied efficiently on this scanner and was omitted.

#### Signal processing

For non-localized (two dimensional) experiments, the first 67 points of the FID in the time dimension were removed to eliminate the distortion from the group delay corresponding to the 13 ms dead time of the Bruker 9.4 T.*^66^* The FID was Fourier transformed and the phase estimated by the entropy minimization method of Chen et al,*^67^*as implemented in MatNMR.**^68^** The baseline was estimated by a modification of the Dietrich first derivative method to generate a binary mask of baseline points,*^69^* followed by spline interpolation using the Whittaker smoother*^70^* to generate a smooth baseline curve.*^71^* The final correction adjusts for the limited number of points in the frequency dimension by continuation of the FID by linear prediction. The 189 points of the FID remaining after truncation in the first step were extrapolated to 1024 points using the “forward-backward” linear prediction method of Zhu and Bax.*^72^* Fourier transforming the FID of the transients from each voxel individually generated the final spectrum. The method proved difficult to apply to the chemical shift imaging experiments and no preprocessing was applied. Spectra for chemical shift imaging experiments are shown in magnitude mode.

#### Low Rank Reconstruction

For the two-dimensional signal matrices generated by non-localized pulse acquire experiments, the rank reduced signal was generated by truncating the SVD by setting the *N-r* diagonal values of the singular value matrix *S* to 0, where *N* is the number of rows in *S* and *r* is the predicted rank. The predicted rank was set to 5 unless otherwise specified, which is equal to the number of independent species in the pyruvate DNP experiment. For the four-dimensional imaging experiments, three methods were tried as described in Figure 7. (1) Applying truncated SVD to each voxel independently (2) Unfolding the four-dimensional tensor into a two-dimensional matrix and applying truncated SVD (a simplified version of LORA*^49^*) (3) Factorizing the four-dimensional tensor by tensor decomposition. The predicted rank was set to 10 for voxel-by-voxel SVD and 16 for the matricization technique, which is the smallest rank which does not introduce substantial distortions in the time averaged image (for example, the inappropriate bleed through of the lipid signal into the tumor mass) as measured by a comparison of the reconstructed and raw time averaged signals. Tensor decomposition was achieved through higher order orthogonal iteration*^73^* in the Matlab NWay package.*^74^* A tensor rank of 8 in the temporal and spatial dimensions and 6 in each spatial dimension was used for the glucose and CSI images.

## Supporting Information Figures

**Figure S1.**
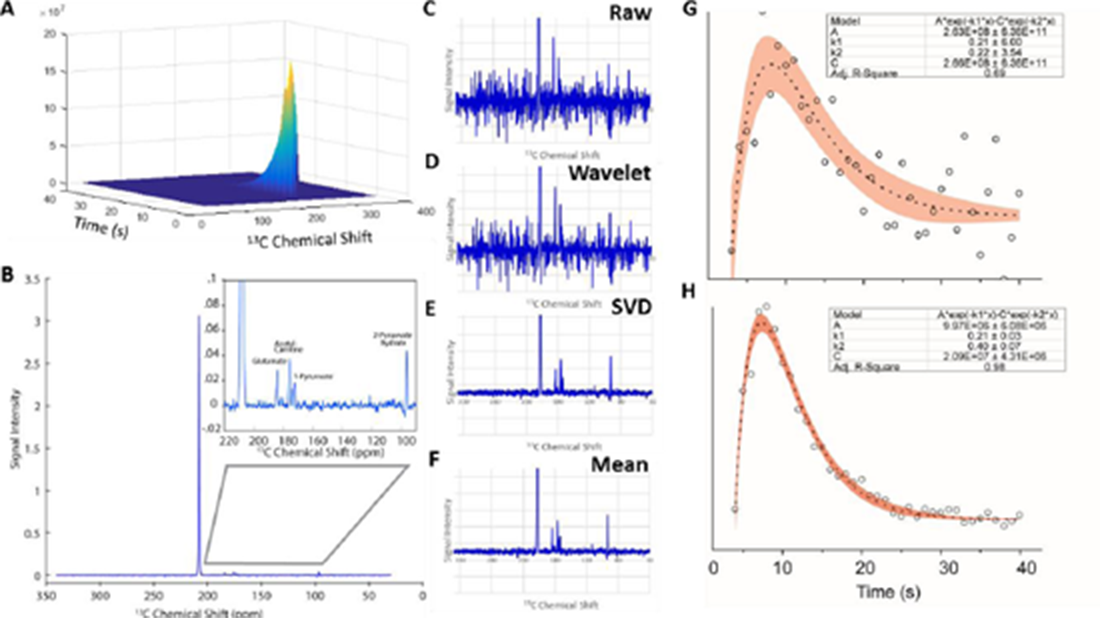
SVD rank reduction improves the precision of kinetic fitting. (A) The raw signal resulting from the brain of an Wistar rat after injection of an 37 mg bolus of pyruvate ^13^C-labeled at the 2 position. **(B)** Spectrum averaged over the 40 s time course. The signal has a very high dynamic range with the metabolites having an intensity only 0.5% of the main 2-pyruvate peak. **(C)** Spectrum 20 seconds after injection. In the raw signal, the metabolites are near the noise level, which is not significantly improved by **(D)** wavelet denoising. By contrast, SVD rank reduction to a rank of 5 results in a significant reduction of noise **(E)** so that the signal resembles the spectra averaged over all time points **(F)**. For glutamate at 184 ppm, the noise in the baseline of the raw signal means the curve fitting is ill-conditioned and the kinetic constants cannot be recovered with any accuracy. **(H)** Improvement in the signal to noise at longer time points by SVD removes the instability and allows reconstruction of the kinetics with greater precision.

**Figure S2.**
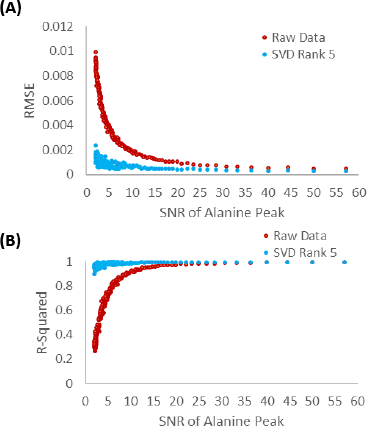
Precision of curve-fitting of the alanine peak in a synthetic data set expressed as **(A)** R-Squared or **(B)** Root Mean Squared Error (RMSE). Use of rank reduction by SVD yielded precise curve-fitting over a large range of signal to noise, while the precision of curve-fitting of the raw data to the biexponential equation *y = A(e^-klt^* e^-K2t^) declined dramatically when the SNR of the metabolite peak became less than 10.

**Figure S3.**
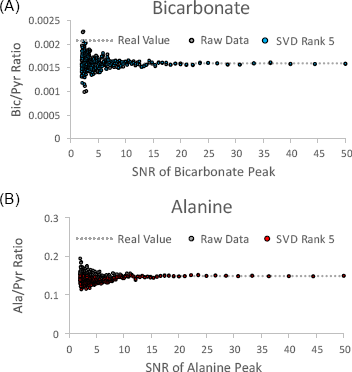
Accuracy of SVD rank reduction from reconstruction of a simulated data set. (A) Bicarbonate-to-pyruvate ratios as a function of the SNR of the bicarbonate peak from the raw and a rank 5 SVD reconstruction (B) Same as above except for the pyruvate peak. A slight bias in the SVD reconstruction exists for the alanine peak at low (<10) signal-to-noise ratios. SVD reconstruction yielded an unbiased estimate at all noise levels for the bicarbonate peak.

**Figure S4.**
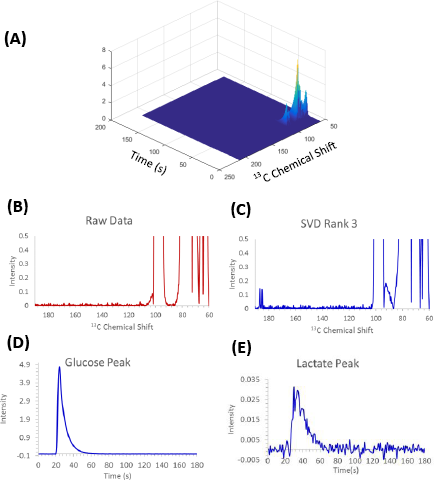
**(A)** Evolution of a hyperpolarized U-C-H glucose tracer after being injected directly into an Hs766t leg xenograft. **(B)** The spectrum at the point of maximum intensity at 24 seconds. Despite the high signal to noise, the only detectable peaks in the spectrum are from glucose with no evidence of metabolic turnover. **(C)** The same spectrum after rank reduction to a rank of 3. A doublet near the expected position of lactate is now apparent. (**D** and **E**). Kinetics of the main glucose and lactate peaks. The lactate peak decays more slowly than the pyruvate peak, reflecting the eventual breakdown of glucose into lactate.

**Figure S5.**
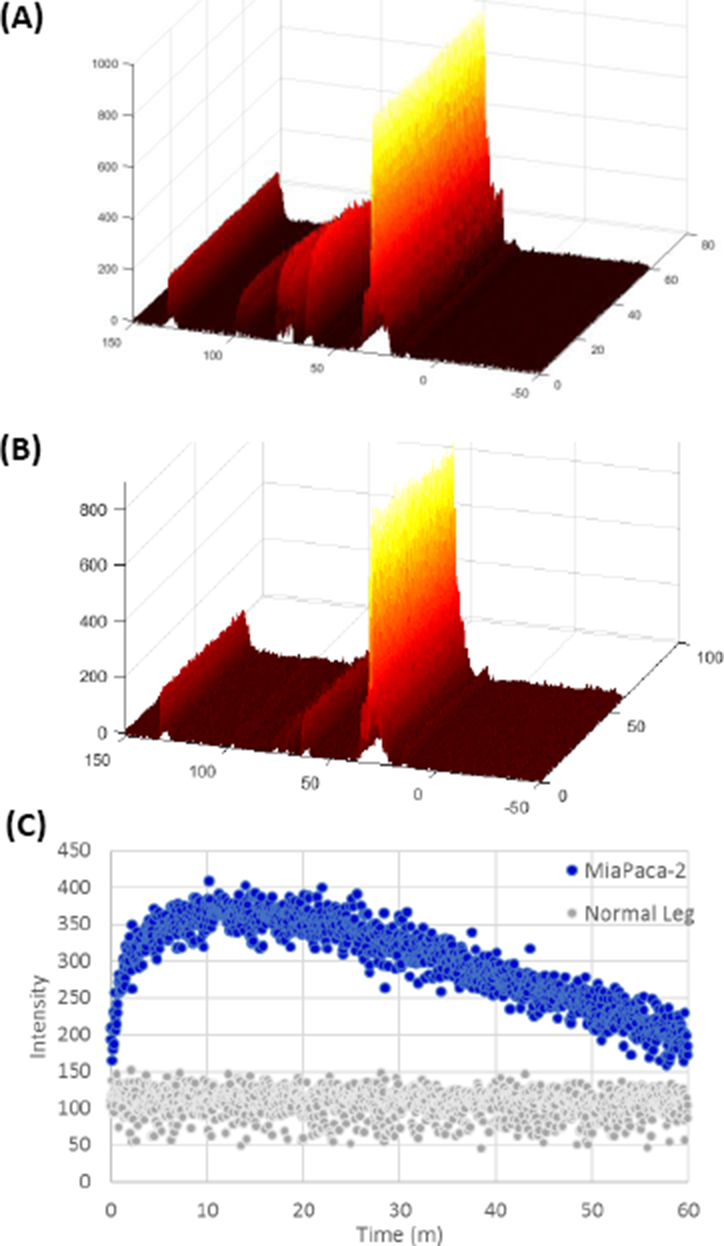
Comparison of Glucose Metabolism. ^13^C signal after injecting 50 mg of uniformly ^13^C labeled glucose into the tail vein of a mouse with a MiaPaca xenograft without hyperpolarization after rank reduction. **(B)** The same experiment on a mouse without a tumor xenograft. **(E)** Kinetics of the signal at 60.5 ppm. No uptake is detectable in the leg without a xenograft, only the constant background signal from lipid glycerol groups.

**Figure S5.**
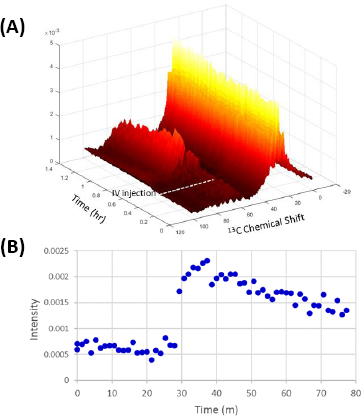
**(A)** Evolution of the C signal at 3T after IV injection of 50 mg U-C glucose. The dashed line indicates the time of injection. **(B)** Kinetics of the glucose signal.

**Figure S6.**
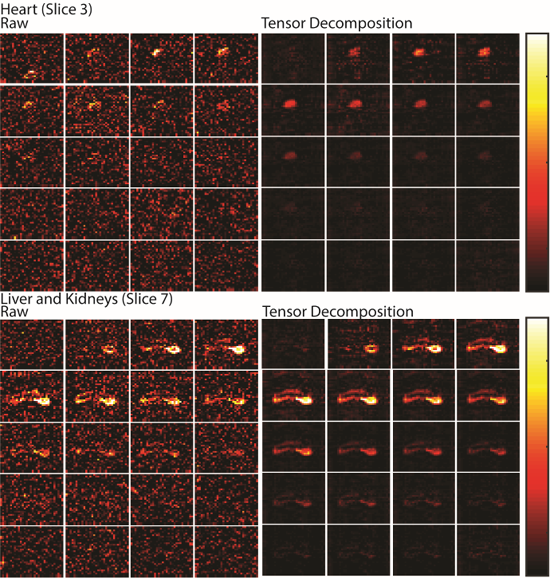
Denoising of Image Data Using Tensor Decomposition. Top Left One slice from a 20 time point image set of lactate production following the injection of hyperpolarized pyruvate centered on the heart. The data was acquired using a spectral selective pulse sequence that yields a series of images for each metabolite, one image for each time point. **Top Right** The same data using the Tensor Decomposition; reducing the rank from [32,32,10,20] to [16,12,8,4]. Bottom The corresponding images from a slice centered on the liver and kidneys. In both cases a roughly 3 fold improvement in signal to noise is observed.

